# A Comprehensive Pangenome Approach to Exploring the Probiotic Potential of *Weissella Confusa*

**DOI:** 10.1101/2024.10.06.616837

**Authors:** Md. Saddam Hossain, Shanjana Rahman Tuli, Nigar Fatima, Sobur Ali, Shovan Basak Moon, Abu Hashem

## Abstract

**Background:** Fermented foods harbour the bacterium *Weissella confusa*, which has probiotic properties but can potentially act as an opportunistic pathogen in humans and animals. Using pangenome analysis, our study aimed to improve the classification and identify functional traits of publicly available *W. confusa* genomes, focusing on evaluating their potential as probiotics.

**Method:** The genomic sequence of 120 strains of *W. confusa* was acquired from the NCBI RefSeq database. The downloaded sequences underwent a quality verification and filtering process. Ultimately, the chosen genomes were examined for comparative genomic analysis. We employed Roary to investigate the pangenome of *W. confusa*. Fisher’s exact test was utilised to analyse contingency tables, with a significance threshold of p < 0.05 (two-tailed).

**Results:** Our investigation revealed that the pangenome of *W. confusa* comprises 1100 core genes, 184 soft-core genes, 1407 shell genes, and 7006 cloud genes. This finding emphasises the “open” aspect of the *W. confusa* pangenome. The comparison of genomes showed that there were no acquired antibiotic resistance genes. However, the strains had different amounts of prophage regions, CRISPR arrays, and plasmids. Our research identified probiotic marker genes (PMGs), with the majority (78%) found in the core and soft-core genomes of various strains of *W. confusa*.

**Conclusions:** An extensive investigation of the *W. confusa* pangenome has concluded that it could be useful as a probiotic. Additional research is necessary to thoroughly evaluate the potential risks.

## Background

Fermented foods, considered among the processed meals consumed by people, boast nutritional and health benefits. They are often referred to as “naturally fortified functional foods” due to their role in promoting healthy gut microbiota, which is crucial for maintaining overall well- being and protection against illnesses (Marco et al., 2017). Over the past decade, researchers have shown a growing interest in fermented foods for their abundance of probiotic bacteria (Shangpliang et al., 2017). According to the Food and Agriculture Organization (FAO) and the World Health Organization (WHO), probiotics are live microorganisms that provide health benefits when consumed in sufficient quantities. Lactic acid bacteria (LAB) are a type of probiotic known for their friendly characteristics their ability to thrive in the gut and their production of various bioactive compounds (peptides, exopolysaccharides (EPS), amylase, protease, lipase enzymes, organic acids) (Mathur et al., 2020). LAB exhibits bio-functional properties such as anti-inflammatory, anti-oxidant, anti-cancer effects, and resilience against stress due to its versatile capabilities (Ringø et al., 2018).

Recently, the species *Weissella* was added to the family of lactic acid bacteria (LAB). There are more than thirty species of *Weissella (Parte, 2018)*. They have been found in food, human, soil, plants, animals, and fish, among other settings. Among them, several strains of *W. confusa* were reported to be opportunistic pathogens and for that reason, there is always a concern about possible infection in the predisposing groups and fortunately, those infections are therapeutically treatable (Ahmed et al., 2022). *Weissella confusa* has been recommended as a potential probiotic due to its desirable phenotypes of high EPS generation, adherence capability, sensitivity to antibiotics, and resistance to low pH, bile salt, and heat (Adebayo- Tayo et al., 2018; Dey et al., 2019; Fusco et al., 2015; K. W. Lee et al., 2012). Furthermore, reports demonstrated that the *W. confusa* strain could utilise phytic acid and create phytase, which decreased the amount of anti-nutritional compounds (Rizzello et al., 2019). Moreover, The German Committee for Biological Agents has classified it as Risk category 1 microorganisms which means they are not associated with disease in healthy adults. Additionally, the American Biological Safety Association (ABSA) has not explicitly assigned it to a risk category. The American Type Culture Collection (ATCC) recommends using the strain ATCC 10881TM at the biosafety level, which makes it unlikely to cause disease in healthy individuals (Ahmed et al., 2022). Despite these promising bio functionalities, *W. confusa* strains have not yet been utilised by producers in the food industry as probiotics, mainly due to the lack of safety verification guidelines that classify disease-causing opportunist vs. desirable probiotics.

The rapid advancement of sequencing methodologies has enabled the sequencing of the genomes of various bacterial strains. This development fueled research on comparative genomics, which investigated the effects of isolation sources on genetic variations, revealed horizontal gene transfer, conducted taxonomic revision, and examined the connections between genotypes and phenotypes (Mao et al., 2021; Singh et al., 2021). These studies illustrated the relationships between genotypes and phenotypes (Wang et al., 2018). Since the recognition of the significance of the *Weissella* species, efforts have been made to investigate their accurate evolutionary connection and genetic properties. Comparative genomic analyses of some *Weissella* species, such as *Weissella hellenica, Weissella cibaria*, and *Weissella paramesenteroides* have been conducted (Li et al., 2017; Panthee et al., 2019; Wan et al., 2023). In one study, novel molecular targets were extracted to identify 11 *Weissella* spp. in food, employing 50 genomes. Moreover, the popular web-based databases have a meagre number of genomes; for instance, Propan contains only 27 genomes, and PanX has no data for *W. confusa*. By 2024, the National Center for Biotechnology Information (NCBI) had uploaded 120 genome assemblies of *W. confusa* strains as RefSeq to its public database. Through comparative genomic analysis, we will be able to understand the capacities and functions of these *W. confusa* strains and learn more about their mode of action, biodiversity, and probiotic gene markers which will help to select a new generation of probiotics and ultimately will enable the potential application of *W. confusa*.

## Methods

### Genome Retrieval

We searched the National Center for Biotechnology Information (NCBI) genome assembly website (https://www.ncbi.nlm.nih.gov/assembly) on 07 February 2024 using the phrase ‘*Weissella confusa*’ (Sayers et al., 2024). We specifically chose the latest RefSeq as the progress bar and downloaded only the DNA data in FASTA format. We utilised the NCBI datasets and dataformat command line interface (CLI) to retrieve the metadata associated with the genomes under investigation.

### Quality Control and Filtering of Downloaded Genomes

The downloaded genomes were assessed for contamination using checkM (Parks et al., 2015), ContEst16S (I. Lee et al., 2017), and BUSCO version 5.6.1 (Simão et al., 2015). To ensure high-quality bacterial genome data for analysis, we implemented a filtering process based on four key metrics: CheckM, ContEst16S, BUSCO, and the number of contigs. We assessed each parameter and assigned a score of either 0 or 1. Genomes with a contamination level greater than 3%, ContEst16S study contamination, BUSCO analysis below 96%, or contigs over 200 were assigned a score of 0 (zero). We selected only genomes with a cumulative score of 3 or higher for further curation. Following the first curation, we utilised the pyani version 0.2.12, a Python package and the TYGS service to compute the average nucleotide identity (ANI) and digital DNA:DNA hybridization (dDDH) values, respectively (Meier-Kolthoff & Göker, 2019; Pritchard et al., 2019). Additionally, we used OrthoFinder v2.5.5 with the default settings to determine the evolutionary relationship between the strains of *W. confusa* utilising the Prokka annotated protein sequences (Emms & Kelly, 2019).

### Comparative Genomics

The Prokaryotic Genome Annotation System (Prokka), version 1.14.5, was used to annotate the whole genome sequences of *W. confusa (Seemann, 2014)*. Subsequently, functional annotation was carried out using Rapid Annotation Subsystems Technology (RAST) (Brettin et al., 2015). The standalone version of ResFinder was used to predict acquired antimicrobial resistance genes, while dbCAN3 was used to predict cazyme genes (Bortolaia et al., 2020; Zheng et al., 2023). PHASTEST (PHAge Search Tool with Enhanced Sequence Translation), a web server, was utilised to thoroughly examine the genome of all strains to identify DNA areas that designate prophages (Wishart et al., 2023). The CRISPR-Cas system, comprised of Clustered Regularly Interspaced Short Palindromic Repeats (CRISPR) and its associated Cas protein, underwent analysis utilising the CRISPRCasFinder (Couvin et al., 2018). The information regarding the presence of plasmids was obtained from the NCBI Assembly page dedicated to each individual strain. The COGclassifier was utilised to categorise prokaryote protein sequences into functional categories based on the COG (Cluster of Orthologous Genes) database (Galperin et al., 2021).

### Pangenome Analysis

The pangenome analysis software Roary v 3.20 was used to calculate the count of core and auxiliary genes in the genomes of *W. confusa (Page et al*., *2015)*. Polymorphic sites were obtained from the Roary output file by utilising SNP-sites (Page et al., 2016). The phylogenetic tree was generated using IQ tree v 1.6.12, and visualised using iTOL v6 (T. Zhou et al., 2023). The eggNOG-mapper v2 was used to predict clustered ortholog groups (COG) for core genes, accessory genes (including soft-core and shell genes), and cloud genes (Cantalapiedra et al., 2021). We conducted Fischer’s exact test to determine the association between the gene frequency groupings (core, accessory, and clouds) and the COG functional category. The significance level was established using the Family-wise Error Rate (FWER) criterion, with a threshold of less than 0.05 after applying the Bonferroni adjustment, or a p-value of less than 0.00092 (Rajput et al., 2023).

## Results

### Data retrieval and Data curation

As of February 19, 2023, a total of 120 assembled genome datasets were available and subsequently downloaded from the NCBI RefSeq database. The total number of assembly levels was 6 for complete assemblies, 19 for scaffold assemblies, and 95 for contig assemblies. However, no reference genome was available, except for one representative genome with the accession number GCF_004771075.1 (**Table S1**). Following the initial curation analysis, it was concluded that three accessions (GCF_018390755.1, GCF_025127155.1, and GCF_025127175.1) exhibited significant contamination. Consequently, genomic data associated with these accessions was removed. In the end, our dataset comprised 117 genomes (**Table S2**).

The mean total size of the genomes and the proportion of GC content in the 117 *W. confusa* strains were 2.27 Mb and 44.69%, respectively. Our datasets had no significant correlation between the total sequence length and the number of contigs or N50 values. The correlation coefficient for the number of contigs and total sequence length was -0.004982388, while the coefficient of determination was 2.482419×10^−05^. Similarly, the correlation coefficient for the contig N50 and total sequence length was 0.3956669, with a coefficient of determination of 0.1565523. The further examination of 117 *W. confusa* genomes using Orthofinder, as well as the study of ANI and dDDH values, did not reveal any instances of incorrectly classified strains within the species (**Tables S3-S4)**. Therefore, all 117 genomes in the data set were included in the ensuing analyses.

### Comparative Genome Analysis

The ResFinder analysis detected no acquired antibiotic resistance genes in the *W. confusa* genomes. A substantial number of genes were identified and categorised in the RAST carbohydrate subsystem and COGclassifier study (**Tables S5-S6 and Fig. 3**). No prophage region was detected in the genomes of LMG 14040 (GCF_016465845.1), BCC 4255 (GCF_016466205.1), and BCC 2330 (GCF_016466355.1). For other individuals, a minimum of one prophage area was identified (**Table S7**). All strains, save for three, contained a single CRISPR array. However, none of the strains in **Table S8** possess the Cas cluster protein. The range of CRISPR arrays observed was from four to one, with strains BCC 4194, BCC 2384, and AM110-142 having the maximum number of CRISPR arrays, each containing four arrays (N = 4). In contrast, three strains, namely Wcp3a, UC 724F13, and 1523, did not exhibit any CRISPR arrays. Plasmid was detected in seven strains, specifically MBF8-1, VTT E-133279, N17, LM1, WiKim51, CYLB30, and VTT E-90392. A total of 16 plasmids were identified in these six strains. The strain CYLB30 included the six plasmids with the highest abundance, as determined in the genome sequence. The mean size of the genome was 15.2 kilobases (kb). (**Table S9**).

**Fig. 1.**
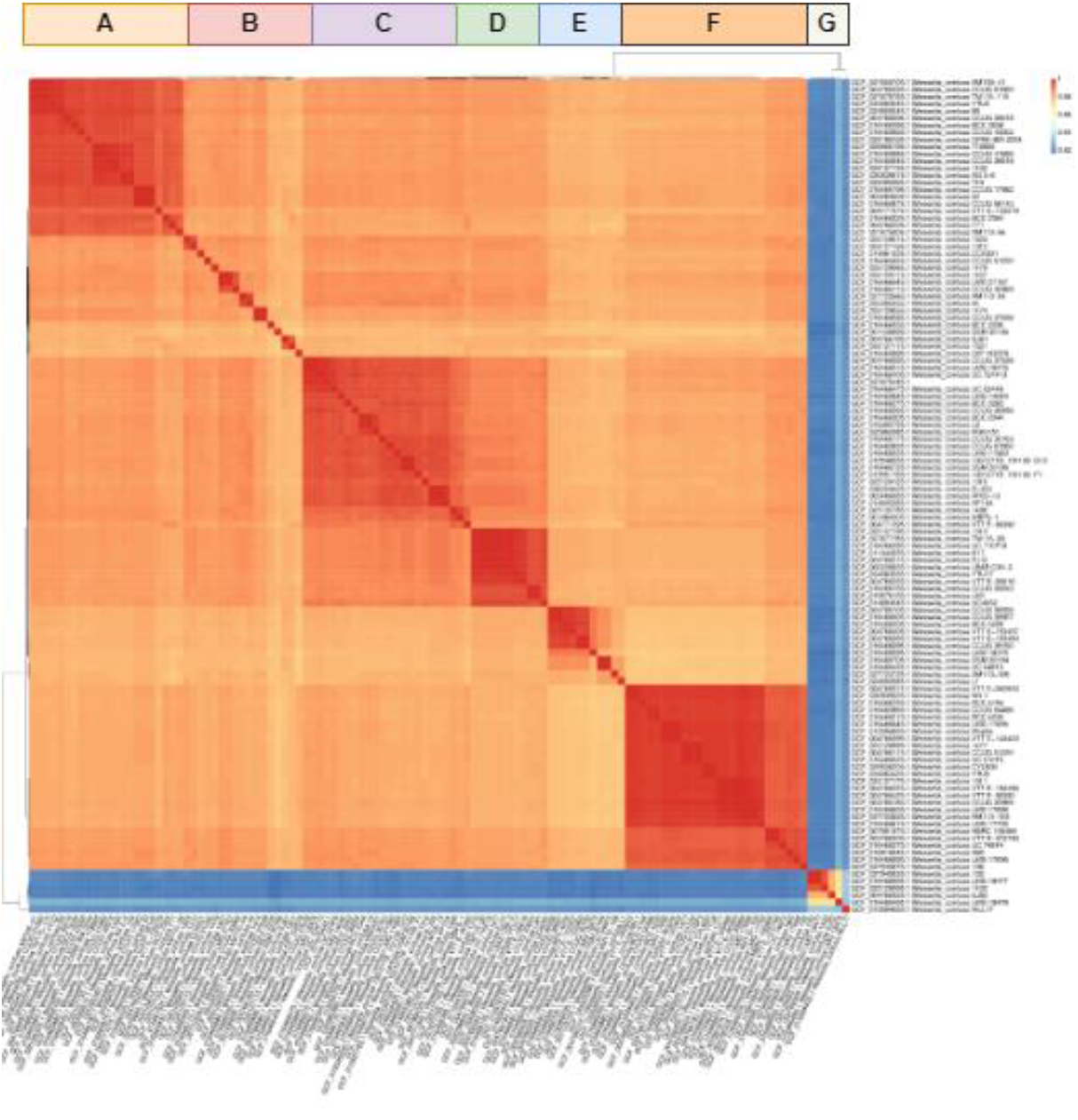
A cluster of *W. confusa* genomes based on average nucleotide identity (ANI) values The six strains (VTT E-062653, VTT E-90392, LMG 718955, LMG 11983, DSM 20196, and TM115-115) clustered apart from the rest of the data set, according to analysis using both pyANI and orthofinder (**Fig. 1-2**). It appears that these strains lacked any specific genetic characteristic that would lead one to classify them as outliers or incorrectly classify them as *W. confusa* strains.

**Fig. 2.**
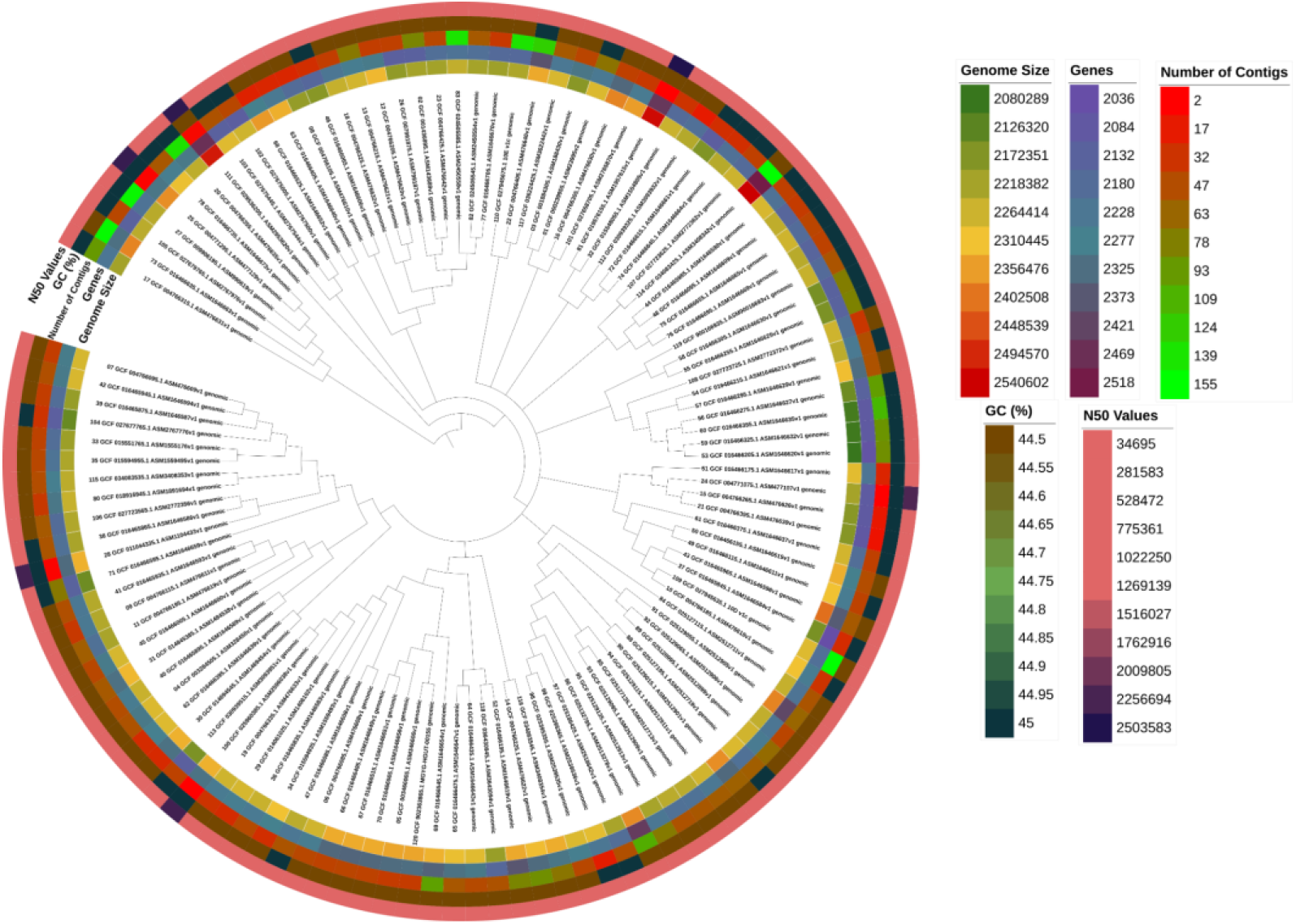
Phylogenetic tree depicting 117 strains of *W. confusa*, constructed using orthologous genes identified with Orthofinder. The accompanying heatmap visualises key genomic characteristics, including genome size, gene count, contig number, GC content, and N50 values for each strain

**Fig. 3.**
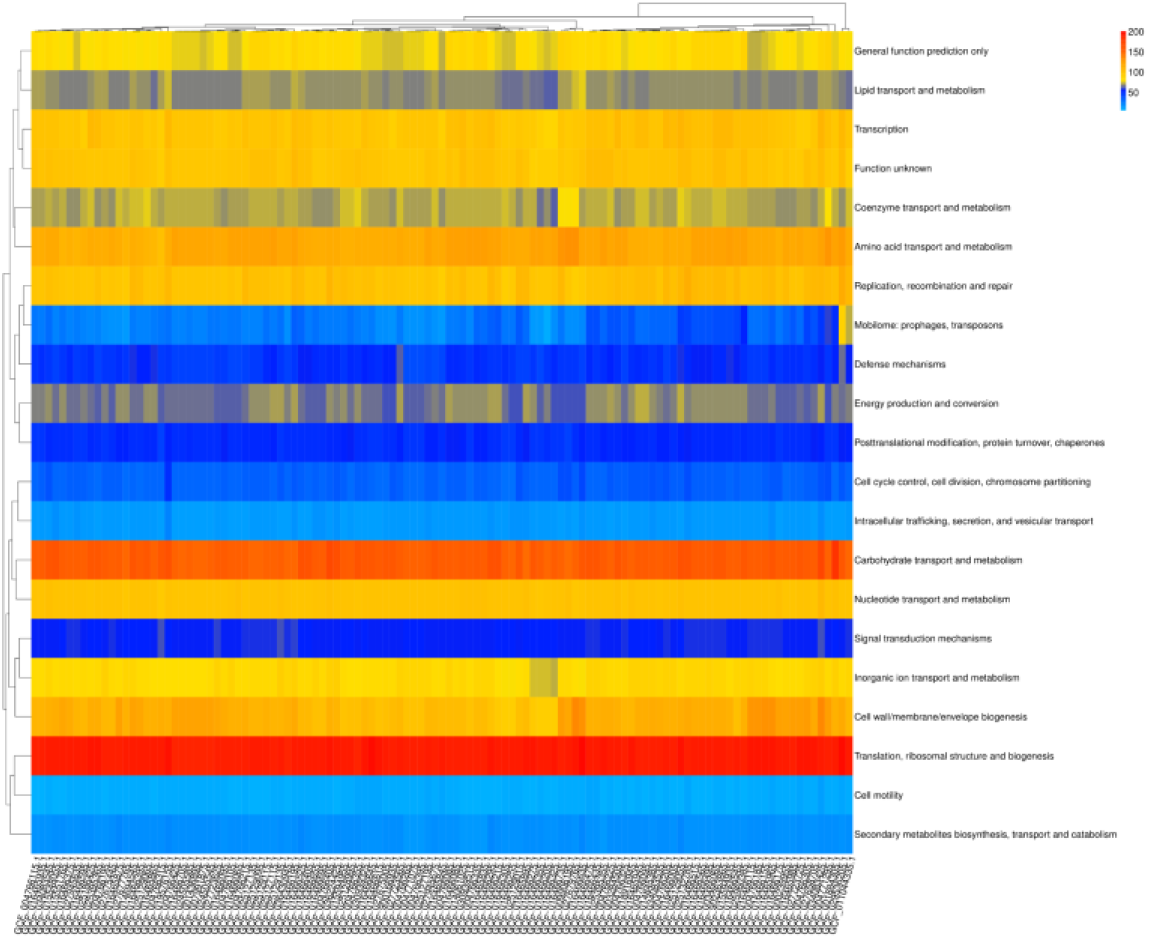
A heatmap of COGs abundances in the 117 *Weissella confus*a genomes

The analysis of CAZymes demonstrated that all genomes possessed genes in the four CAZymes gene families (CBM, CE, GH, and GT), and the strains exhibited functional similarities. Among all the strains, only strain 1001271B_151109_G12 (GCF_015548055.1) exhibited an extra PL family, as depicted in **Fig. 4** and **Table S10**. The enzymes GH33, 16, 29, 95, 20, 2, 35, 42, 98, 101, 129, 89, 85, and 84 are known to participate in breaking down mucin (Glover et al., 2022). *Weissella confusa* MBF8-1 and *Weissella confusa* CIRM-BIA 2234 lack the glycosyl hydrolases necessary for mucin degradation, while others possess only GH2 in a single copy.

**Fig. 4.**
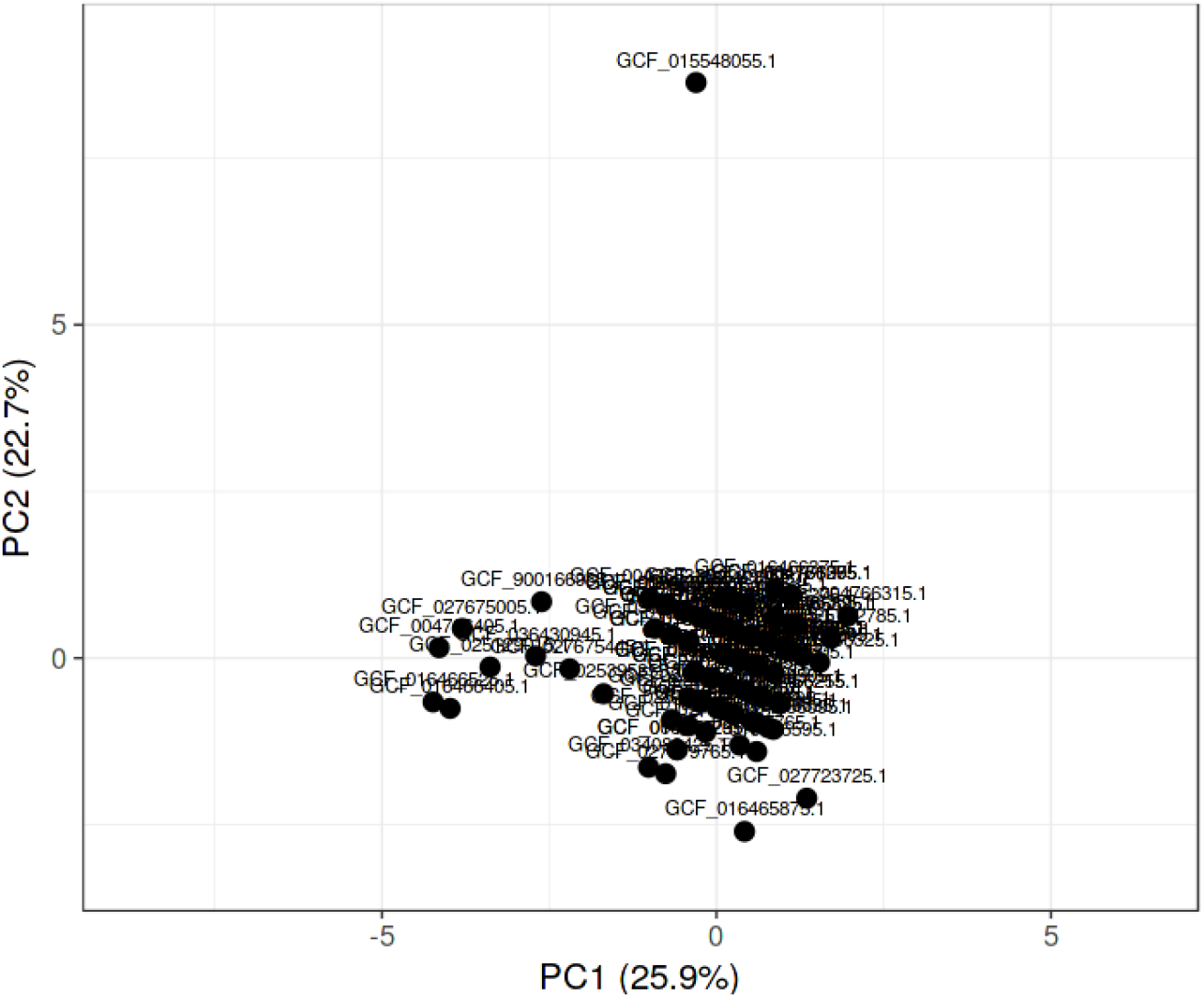
Principal component analysis (PCA) on *W. confusa* strains using predicted cazymes. The X and Y axes represent principal component 1 and principal component 2, respectively, explaining 25.9% and 22.7% of the total variance. The dataset comprises 117 data points.

### Pangenome analysis of *Weissella confusa*

The investigation of the *W. confusa* pangenome using Roary revealed a total of 9697 genes. Among them, there were 1100 core genes, 184 soft core genes, 1407 shell genes, and 7006 cloud genes (**Fig. 5a**). The presence of a significant number of cloud genes indicates that there is considerable diversity among the 117 *W. confusa* strains under consideration, which emphasises the expansive nature of the *W. confusa* pan-genome (**Fig. 5b**). However, we observed a gradual decrease in the number of new genes in relation to the number of genomes analysed (**Fig. 5c**). A matrix based on the presence and absence of genes was also generated (**Fig. 5d**)

**Fig. 5.**
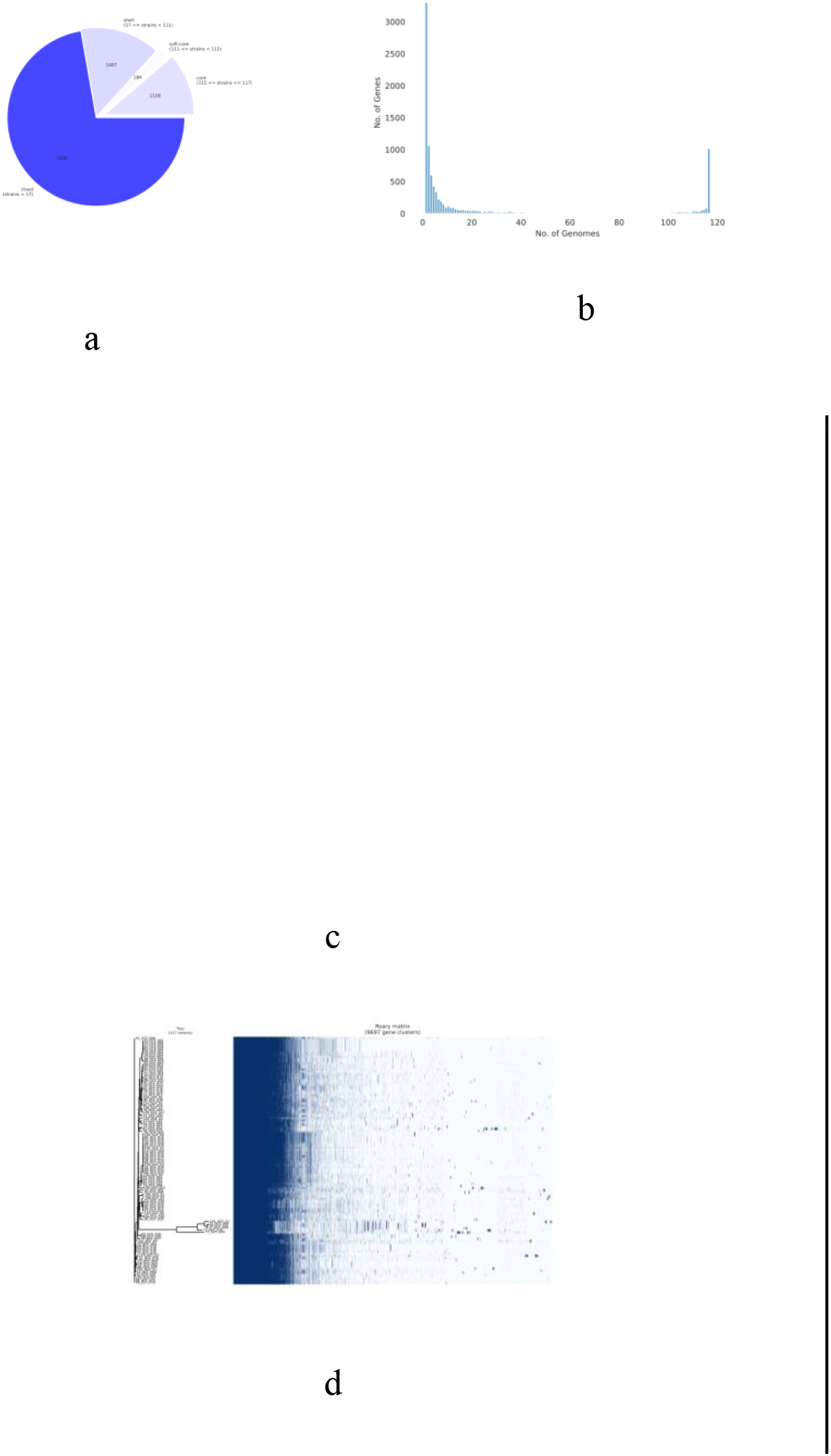
*Weissella confusa* pan-genome. (a) A pie chart illustrates the distribution of genes among the core, soft core, shell, and cloud categories in the *W. confusa* species; (b) A graph illustrating the relationship between the frequency of genes and the number of genomes; (c) A visualisation of *W. confusa* gene content as the pangenome changes with the random inclusion of genomes for analysis; (d) The matrix of the accessory and core genes of the isolates is represented by blue (present) and white (absent) fragments.

A phylotree was constructed using single nucleotide polymorphisms (SNPs), which revealed that all strains originated from a common ancestor. The SNP-based, core-genome phylogenetic tree validated the distinctiveness of the six strains, which pyANI had already detected. None of the remaining strains in the clades were consistently found in both phylogenetic trees, which were constructed using orthologous genes and SNPs. Furthermore, there was a significant difference in strain distribution among the clades (p = 0.00133). Additionally, the phylogenetic tree based on single nucleotide polymorphisms (SNPs) showed a significant difference when stratified by isolation source category (p = 0.0454, Food vs Human) (**Fig. 6**).

**Fig. 6.**
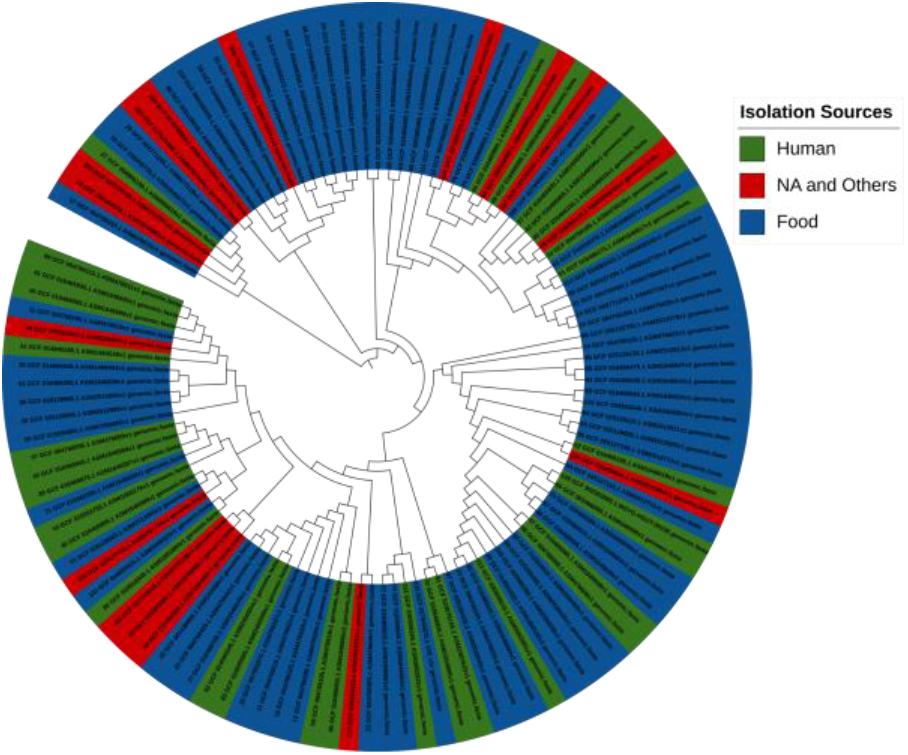
Phylogenetic tree of the 117 *W. confusa* strains. Tree based on single-nucleotide polymorphisms (SNPs) identified by SNP-sites among the strains. Different colours show isolation sources. (NA= Not applicable)

**Fig. 7:**
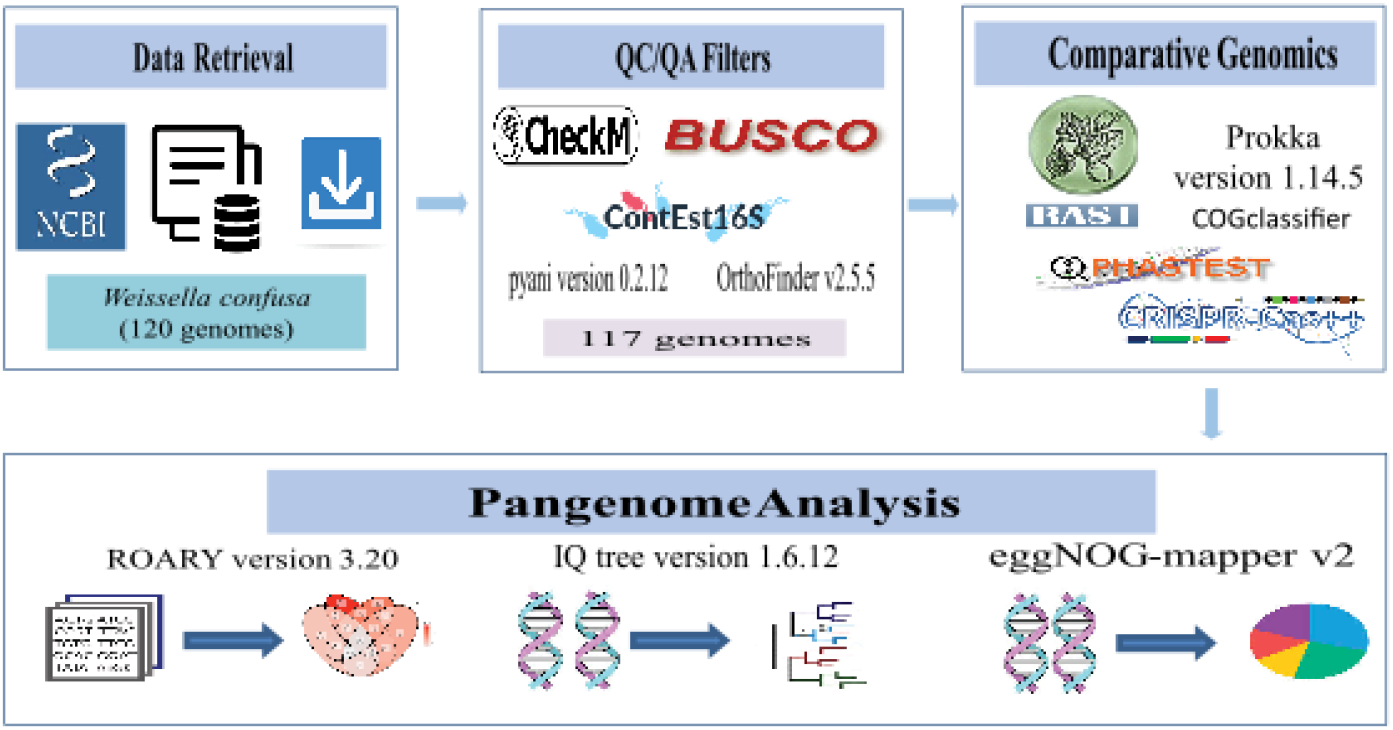
A graphical representation of the workflow

Additionally, a Fisher’s exact test was conducted to validate the statistical enrichment of the projected functions assigned by eggNOG among the genomes categorised as Core, Accessory, and Clouds. (**Table S12**). Out of the 26 COG categories, 7 showed statistical significance and were shown to be enriched in the core genome. The categories included translation, ribosomal structure and biogenesis (J), signal transduction mechanisms (T), post-translational modification, protein turnover, chaperones (O), nucleotide transport and metabolism (F), coenzyme transport and metabolism (H), lipid transport and metabolism (I), and inorganic ion transport and metabolism (P). Regarding the accessory genome, we have identified three COG functional categories that were statistically significant and enriched. These categories include carbohydrate transport and metabolism (G), inorganic ion transport and metabolism (P), and function unknown (S). Three COG categories that showed statistical enrichment were observed in the cloud genome case. The three main biological processes involved were replication, recombination, and repair (L), defensive mechanisms (V), and cell wall/membrane/envelope biogenesis (M).

### ‘Probiotic marker genes’ in the *Weissella confusa* pangenome

A probiotic bacterium must possess the capacity to survive and temporarily remain in the gastrointestinal tract to exert a favourable impact. In addition to mobile genetic elements (MGEs) and genes related to the adaptive immune system, the presence of genes that enable resistance to host stressful situations, as well as the ability to break down bile salts, is a crucial factor in identifying the important genes that define a bacteria as a possible probiotic.

In this work, we compiled a list of ‘probiotic marker genes’ (PMGs) that are important for stress tolerance (acid, osmotic, oxidative, temperature), bile salt hydrolase activity, adhesion ability, and gut persistence (**Table S11**). We aimed to identify any distinctive characteristics shared by all the *W. confusa* strains included in our research. The report has included the annotation and the presence or absence status for 60 PMGs. Based on the comparative pan-genome study conducted with Roary, we found that around 78% of the analysed PMGs were part of the core/soft-core genome, which included strains ranging from 95% to 100% similarity. The bshA gene, associated with tolerating bile salts, was found exclusively in the *W. confusa* 17 strains. The gene xylA, which is implicated in intestinal persistence, was found exclusively in three strains. Two strains were identified to contain the PMG clpP1_2, which is involved in acid stress (Carpi et al., 2022).

## Discussion

Recent studies have shown that analysing the pangenomes of specific species provides valuable information that has yet to be fully understood (Rajput et al., 2023). This study is the first to identify the pangenome of *W. confusa*, a promising candidate for probiotics, by analysing a large number of genomes (N = 117) from the NCBI RefSeq database. To be more specific, we wanted to find out what probiotic properties each strain had by looking at their plasmid content, mobile genetic elements (MGEs), adaptive immune system, and probiotic marker genes (PMGs). Additionally, we categorised the *W. confusa* pangenome into four gene categories (‘core’, ‘soft core’, ‘shell’, and ‘cloud’) to facilitate genetic engineering techniques to reduce and optimise the genome.

To avoid significant biases that could affect the interpretation and analysis of the results, we primarily concentrated on carefully selected genomic data of *W. confusa* strains. This would open the door to a more in-depth analysis of this bacterium’s genetic makeup.

Prophage regions and CRISPR loci may enhance a bacterial strain’s ability to adapt to its surroundings (Peters et al., 2017). The majority of the strains in this investigation (97%) had bacteriophage sequences and CRISPR arrays. Seven of the strains under investigation also contained plasmids. Since mobile genetic elements (MGEs) play a major role in antibiotic resistance, identifying and characterising MGEs is crucial to determining the probiotic potential of a particular strain.

RAST annotation and COGclassifier analysis revealed that the metabolism of carbohydrates enriched a significant portion of the *W. confusa* genomes. The CAZymes study also found that glycoside hydrolases (GHs), glycoside transferases (GTs), carbohydrate-binding modules (CBMs), and carbohydrate esterases (CEs) were the most common enzymes in the *W. confusa* genomes. The presence of abundant GHs in *W. confusa* raises the possibility that it could function similarly if used as a probiotic supplement (Al-Emran et al., 2022). The degradation of mucin glycans requires the synchronised activity of several glycosyl hydrolases produced by bacteria with the ability to break down mucin (Glover et al., 2022). Our study demonstrated that no strain of *W. confusa* contains several glycosyl hydrolases known to participate in breaking down mucin. These results suggest that *W. confusa* strains are probably harmless and do not penetrate the mucosal barrier since they are incapable of breaking down gastrointestinal mucin (J. S. Zhou et al., 2001). Our study’s weakness is that our dataset only included six complete genomes.

However, the comparative genomic analysis of the study gives new insights into the genetic diversity and richness of *W. confusa*. This highlights the need to screen new strains because bacterial genomes constantly change. By looking at the core, auxiliary, and unique genes in genomes, it is possible to tell the difference between strains with different traits, which opens the door to finding good probiotic candidates.

## Conclusions

To summarise, the comprehensive study of the *W. confusa* pangenome has demonstrated its potential contribution to maintaining gut health. More research needs to be done on the organism’s functional characterization and safety assessment to fully use its untapped potential as a probiotic agent. Unlocking its full potential could lead to the development of new probiotic products that enhance gastrointestinal health.

## Declarations

## Ethics approval and consent to participate

Not applicable

## Consent for publication

Not applicable

## Availability of data and materials

The data provided in this work can be obtained upon a reasonable request from Dr. Abu Hashem, the corresponding author.

## Authors’ contributions

MSH and AH: conceptualization, study design, manuscript writing, and submission; SRT, NF & SA: experimental investigation; MSH and SBM: bioinformatic research; AH: study guidance and final review.

## Acknowledgments

Not applicable

## Authors’ information

Md. Saddam Hossain, Shanjana Rahman Tuli, and Nigar Fatima contributed equally to this work.

## Graphical abstract

